## Supplemental material for "Hearables: Feasibility of Recording Cardiac Rhythms from Single Ear Locations"

*Artifact rejection: Experiment B.*

Large amplitude deflections were removed from each ear ECG signal via a threshold rejection procedure. The threshold was set to 30µV and 200µV, respectively, for the single ear and cross-ear signals. Blink artifact removal consisted of subtracting the grand-average blink artifact from each of the ear ECG signals at each instance of a blink event in the VEOG signal. The grand-average blink was only removed if the Pearson correlation with the time-aligned ear ECG signal exceeded r = 0.6. Before subtraction, the grand-average blink was scaled by a factor equal to RMS_ECG/RMS_blink, where RMS_ECG denotes the root-mean-square amplitude of the segment of the ear ECG signal in question, and RMS_blink denotes the root-mean-square amplitude of the grand-average blink signal from that same channel. Segments were 400ms in length and centred around the peak of the blink event in the VEOG signal. Blink events in the VEOG signal were found using a peak detection algorithm ‘findpeaks’ in Matlab, with manually set peak-height input parameters for each subject. Spectral coherence between the ear ECG and VEOG signals was reduced by a mean of 0.2, 0.3, and 0.1 for the left ear, right ear, and cross ear ECG signals after the described procedure [1]. The function ‘mscohere’ in Matlab was used to calculate the spectral coherence. These moderate decreases in spectral coherence are expected as a result of the small amplitude of blink artifacts in the ear ECG signal [2].

*References*

[1] Kay, Steven M. Modern Spectral Estimation. Englewood Cliffs, NJ: Prentice-Hall, 1988.

[2] Yarici, Metin, Mike Thornton, and Danilo Mandic. "Ear-EEG Sensitivity Modelling for Neural Sources and Ocular Artifacts." *Frontiers in Neuroscience,* (2022) [in press]
